# Conserved patterns of alternative splicing in response to cold acclimation in fish

**DOI:** 10.1101/429704

**Authors:** Timothy M. Healy, Patricia M. Schulte

**Affiliations:** The University of British Columbia, Department of Zoology, 6270 University Boulevard, Vancouver, British Columbia, Canada, V6T 1Z4

**Keywords:** temperature, differential exon usage, phenotypic plasticity, killifish, stickleback, zebrafish

## Abstract

Phenotypic plasticity is an important aspect of an organism’s response to environmental change that often requires the modulation of gene expression. These changes in gene expression can be quantitative as a result of increases or decreases in the amounts of specific transcripts, or qualitative as a result of the expression of alternative transcripts from the same gene (e.g., via alternative splicing of pre-mRNAs). Although the role of quantitative changes in gene expression in phenotypic plasticity is well known, relatively few studies have examined the role of qualitative changes. Here, we use skeletal muscle RNA-seq data from Atlantic killifish (*Fundulus heteroclitus*), threespine stickleback (*Gasterosteus aculeatus*) and zebrafish (*Danio rerio*) to investigate the extent of qualitative changes in gene expression in response to cold. Fewer genes demonstrated alternative splicing than differential expression as a result of cold acclimation; however, differences in splicing were detected for between 426 and 866 genes depending on species, indicating that large numbers of qualitative changes in gene expression are associated with cold acclimation. Many of these alternatively spliced genes were also differentially expressed, and there was functional enrichment for involvement in muscle contraction among the genes demonstrating qualitative changes in response to cold acclimation. Additionally, there was a common group of 29 genes with cold-acclimation-mediated changes in splicing in all three species, suggesting that there may be a conserved set of genes with expression patterns that respond qualitatively to prolonged cold temperatures across fishes.

**Summary statement:** Qualitative changes in gene expression, such as those mediated by alternative splicing of mRNAs, are involved in phenotypic plasticity in response to prolonged cold acclimation in ectothermic animals

## Introduction

Phenotypic plasticity, which is the ability of organisms to express environmentally mediated alternative phenotypes without genetic change (e.g., Travis, 1994; West-Eberhard, 2003), plays a critical role in determining organismal responses to a changing environment (e.g., Schulte et al., 2011). For example, reversible metabolic plasticity in ectotherms has the potential to increase resilience to anthropogenic climate change (Seebacher et al., 2015). One of the primary mechanisms underlying this phenotypic plasticity is thought to be changes in gene expression (Schlichting and Pigliucci, 1993; Schlichting and Smith, 2002), which can occur in one of two general ways: (1) quantitative changes in which the transcripts of a gene increase or decrease in number in the cell, or (2) qualitative changes in which the nature of the transcripts being expressed changes as a result of mechanisms such as alternative mRNA splicing or RNA editing (Schlichting and Pigliucci, 1993; Gerber and Keller, 2001; Hochachka et al., 2001; Schlichting and Smith, 2002; Schulte, 2004; Rosenthal, 2015). Quantitative changes in gene expression in response to environmental change have been widely studied (e.g., Gracey et al., 2004; Buckley et al., 2006; Garcia et al., 2012; Moya et al., 2012; Schoville et al., 2012; Zhao et al., 2012; Mandic et al., 2014), whereas, despite previous suggestions that qualitative changes are likely also an important mechanism underlying plasticity (Hochachka et al., 2001; Schulte, 2004), surprisingly little is known about qualitative mRNA responses to environmental stressors in animals (Somero, 2018).

Some evidence for the potential importance of qualitative changes in gene expression comes from studies of the phenomenon of RNA editing, in which there is enzymatic conversion of individual bases within a transcript that result in changes in the amino acid sequence of the subsequently translated protein (Nisihikura, 2016; Montiel-Gonzalez et al., 2016). For example, it has been shown that differences in RNA editing of voltage-gated potassium channels between tropical and polar Octopus species results in channels with functional properties suitable for their different habitats (Garrett and Rosenthal, 2012), suggesting that RNA editing provides a mechanism allowing environmental adaptation. However, the role of RNA editing in plasticity has not been well studied (Rosenthal, 2015), and RNA editing appears to be relatively rare in animal taxa other than cephalopods (Liscovitch-Brauer et al., 2018). In contrast, alternative splicing, a process in which different mature transcripts are produced from a single gene (Modrek and Lee, 2002), is widespread across taxa (Tapial et al., 2017), and thus has the potential to be a common mechanism underlying phenotypic plasticity in many species. Alternative splicing in the response to environmental stress is widely observed in plants (Reddy et al., 2013; Capovilla et al., 2015; Filichkin et al., 2015; Thatcher et al., 2015; Calixto et al., 2018), but this mechanism has rarely been examined in animals (although see Polley et al., 2003; Huang et al., 2016; Jakšić and Schlötterer, 2016; Hopkins et al., 2018; Tan et al., 2018; Xia et al., 2018), and much remains unknown about its importance.

Plasticity in response to chronic cold temperature is a particularly promising avenue to investigate the potential role of qualitative changes in gene expression in regulating reversible phenotypic change. Alternatively spliced variants of Δ9-acyl CoA desaturase with temperature-sensitive expression patterns have been detected in common carp, *Cyprinus carpio* Linnaeus (Polley et al., 2003), and many transcriptomic studies examining quantitative changes in gene expression as a result of acclimation to low temperature have observed two key findings: (1) there is generally bias for up-regulation of gene expression as a result of cold acclimation, and (2) there are generally functional enrichments for gene expression and RNA splicing in the genes that are differentially expressed in response to chronic cold temperatures (e.g., Gracey et al., 2004; Long et al., 2012, 2013; Scott and Johnston, 2012; Rebl et al., 2013; Bilyk and Cheng, 2014; Mininni et al., 2014; Morris et al., 2014; Liang et al., 2015; Healy et al., 2017; Metzger and Schulte, 2018). Moreover, these enrichments are often the most well supported enrichments statistically, suggesting changes in gene expression and splicing are key aspects of organismal responses when acclimated to low temperatures. However, splicing of mRNA transcripts is a normal component of RNA processing (e.g., Papasaikas and Valcárcel, 2016), and because cold acclimation typically results in up-regulation of gene expression overall, functional enrichment for genes involved in RNA splicing could simply be a consequence of up-regulated gene expression in general without substantial changes in splicing patterns or qualitative mRNA expression. Therefore, although changes in pre-mRNA splicing are promising candidate mechanisms for the basis of phenotypic plasticity, the extent to which splicing patterns vary in response to chronic temperature change in ectothermic animals remains unknown (Somero, 2018).

The lack of studies addressing temperature-mediated qualitative changes in gene expression is somewhat surprising, because data collected by RNA-seq, including those from previous studies, are amenable for the analysis of changes in splicing patterns or differential exon usage, and modifications of standard RNA-seq analysis packages that enable tests for differential exon usage are publically available (e.g., Anders et al., 2012). In the current study, we take advantage of previously published studies in three species of fish: Atlantic killifish (*Fundulus heteroclitus* Linnaeus; Healy et al., 2017), threespine stickleback (*Gasterosteus aculeatus* Linnaeus; Metzger and Schulte, 2018) and zebrafish (*Danio rerio* Hamilton; Scott and Johnston, 2012). We re-analyze the data from these studies with an overall goal of assessing whether changes in mRNA splicing play a key role in plastic responses as a result of thermal acclimation in ectotherms. To address this goal, we focus on the following questions: (1) What is the extent of alternative splicing in response to cold acclimation? (2) Are alternatively spliced (AS) genes also differentially expressed (DE) genes? (3) What are the potential cellular functions that are influenced by qualitative changes in gene expression? (4) Are there conserved patterns of alternative splicing across species?

## Materials and methods

### Data acquisition

Sequencing reads were obtained for three species of fish from previously published studies investigating quantitative changes in gene expression in response to acclimation to low temperatures: *F. heteroclitus* (Healy et al., 2017: National Center for Biotechnology Information [NCBI] Sequence Read Archive [SRA], SRP091735), *G. aculeatus* (Metzger and Schulte, 2018: NCBI SRA, SRP135801), and *D. rerio* (Scott and Johnston, 2012: European Bioinformatics Institute [EBI] Array Express Archive, EMTAB-1155). From each of these studies, we utilized RNA-seq data from skeletal muscle collected from individuals acclimated to typical laboratory holding temperatures (*F. heteroclitus*: 15°C, n = 8; *G. aculeatus*: 18°C, n = 6; *D. rerio*: 27°C, n = 4), and individuals acclimated to relatively cold temperatures (*F. heteroclitus*: 5°C, n = 8; *G*. *aculeatus*: 8°C, n = 6; *D. rerio*: 16°C, n = 4). Thus, the cold acclimation treatments in the current study represent approximately 10°C decreases in acclimation temperature compared to typical laboratory holding temperatures for all three species. *F. heteroclitus* were wild-caught and held under laboratory conditions (15°C) for at least a month prior to the start of experimental acclimations which were 4 weeks long. *G. aculeatus* and *D. rerio* were lab-raised at 18°C and 27°C, respectively, then experimental acclimations in adult individuals were 4 weeks and 30 days, respectively. Holding photoperiods and salinities varied across species (*F. heteroclitus*: 12L:12D, 20 ppt; *G. aculeatus*: 14L:10D, 20 ppt; *D. rerio*: 12L:12D, 0 ppt), but were consistent between acclimation treatments within each species. Additional methodological details for fish handling, muscle sampling and RNA-seq can be found in Healy et al. (2017), Metzger and Schulte (2018) and Scott and Johnston (2012).

### Gene-wise read mapping and analysis of differential expression

To ensure differences in analytical pipelines among Healy et al. (2017), Metzger and Schulte (2018) and Scott and Johnston (2012) did not affect our comparisons of differential expression patterns among species, and differential expression and alternative splicing within species, we re-analyzed differential gene expression as a result of cold acclimation in each species using a common analytical approach. Sequencing reads were mapped to the appropriate reference genome for each species (*F. heteroclitus* reference genome 3.0.2 [NCBI]; *G. aculeatus* reference genome [ENSEMBL v88]; *D. rerio* reference genome [ENSEMBL v88]) with CLC Genomics Workbench v8.5 (CLC bio Qiagen®, Aarhus, Denmark). Unique exon counts were then exported and analyzed for differential expression using R v3.4.0 (R Core Team, 2017) with the *DESeq2* package v1.16.1 (Love et al., 2014). Differential expression was assessed with the approaches of Healy et al. (2017), which follow the recommendations of Lin et al. (2016). In brief, for each species, counts were normalized across libraries using the relative log expression method (Anders and Huber, 2010) with subsets of genes excluding likely DE genes that were detected in preliminary analyses, genes were filtered for low expression using counts per million cutoffs equivalent to 10 counts in the smallest library, and dispersions were calculated using the default methods of *DESeq2*. Healy et al. (2017) used the *edgeR* package (Robinson et al., 2010) rather than *DESeq2*; however, when parallel methodological steps are utilized, these packages generally result in similar estimates of differential gene expression (Lin et al., 2016). In the current study, we chose *DESeq2*, because the *DEXSeq* package (Anders et al., 2012) that we used for the differential exon usage analysis (below) is an extension of *DESeq2*, and our desire was to minimize bias in our comparisons of mRNA expression and alternative splicing due to differences in bioinformatic techniques. Differential expression was tested by likelihood ratio tests (α = 0.05), and false discovery rate was controlled using the Benjamini-Hochberg method (Benjamini and Hochberg, 1995).

### Exon-wise read mapping and analysis of differences in alternative splicing

Differential exon usage was assessed using R and the package *DEXSeq* v1.22.0 (Anders et al., 2012). Note that *DEXSeq* examines all changes in exon usage, which includes differences in both splicing patterns and usage of alternative transcription start sites. In this study, we refer to all significant effects detected with *DEXSeq* as “alternative splicing” to help clearly distinguish these results from those for differential gene expression; however, it is important to acknowledge that some of our results likely involve changes in transcriptional site usage rather than splicing of pre-mRNAs after transcription has been completed. Regardless, all of these effects result in the expression of alternative mRNAs from the same gene. The first step of the *DEXseq* analysis involves preparing a “flattened” general feature format (GFF) file for the genome of each species, which essentially compresses different transcripts produced from the same gene into a single gene model with non-overlapping exon segments. Reads are then mapped to exon segments, and differential exon usage is detected by differences in read ratios for individual exon segments compared to differences in read ratios across all exon segments for a gene. We performed GFF file flattening and read mapping using *python* as described in the *DEXSeq* manual (available at http://bioconductor.org/packages/release/bioc/html/DEXSeq.html). *DEXSeq* has been designed to perform flattening with GFF files in standard ENSEMBL format. Thus, for *G. aculeatus* and *D. rerio* we were able to use the GFF files available for these reference genomes from ENSEMBL. Because the *F. heteroclitus* genome is hosted by NCBI, the GFF file available for this genome does not follow ENSEMBL formatting. Consequently, we converted the *F. heteroclitus* GFF file to the correct format to allow flattening with *DEXSeq*. We have made our converted GFF file for *F. heteroclitus* available online (*the file will be submitted as part of an EBI BioStudy should the manuscript be accepted*), but other than formatting it is essentially equivalent to the publically available GFF file from NCBI.

Exon read counts were analyzed for differential exon usage following the guidelines in the *DEXSeq* manual. Read counts were normalized, dispersions were calculated, and then differential exon usage was tested (α = 0.05) using the default methods for *DEXSeq*. As for our analyses of differential gene expression, corrections for multiple comparisons in our differential exon usage analysis were made using the Benjamini-Hochberg method (Benjamini and Hochberg, 1995).

### Analysis of functional enrichment

Functional enrichment analyses for the sets of genes demonstrating differential gene expression or alternative splicing were conducted with the R package *goseq* v1.28.0 (Young et al., 2010), and species-specific databases of Gene Ontology (GO) terms. GO annotations for the genes in the *D. rerio* reference genome were obtained from ENSEMBL BioMart. *F. heteroclitus* GO terms were determined from UniProt accession numbers for zebrafish, mouse (*Mus musculus*), and human (*Homo sapiens*) orthologs of killifish genes (Reid et al., 2016), as in Healy et al. (2017). *G. aculeatus* gene GO annotations were accessed from the GO database in Metzger and Schulte (2016). All GO databases have been provided online (*the files will be submitted as part of an EBI BioStudy should the manuscript be accepted*). We assessed significant enrichment of GO functional terms for (1) DE and AS genes in each species separately, and (2) common DE or AS genes across the three species. In all cases α was set at 0.05, and *p*-values were corrected to account for false discovery rate using the Benjamini-Hochberg method (Benjamini and Hochberg, 1995).

### Identification of putative orthologs

Comparison of DE and AS genes within each species was straightforward, as genes could be matched based on their gene identifiers. However, to compare DE or AS genes across species, and thus to identify genes that were commonly differentially expressed or alternatively spliced as a result of cold acclimation, it was necessary to identify putative orthologous genes across the species.

A reciprocal basic local alignment search tool (BLAST) strategy was used to identify putative orthologs (Moreno-Hagelsieb and Latimer, 2007). We downloaded the NCBI command-line based BLAST program, and created a local BLAST database for all the mRNA transcripts associated with the reference genomes for each species in our study (3 databases total). For the DE genes in each species, we queried the mRNA databases for the other two species. We kept the top 10 BLAST hits for each gene, and then filtered hits for e-value < 0.000001 and query coverage ≥ 33.3%. For some genes, several of the top BLAST results with e-value < 0.000001 mapped to different regions of the same target transcript. In these cases, the non-overlapping query coverage was summed across results prior to application of the coverage filter. This procedure was repeated for the lists of genes demonstrating alternatively splicing, and resulted in high-quality BLAST results for between 72.1% and 96.0% of the genes for each pair of species across the DE or AS genes. Putative orthologs were then determined by limiting BLAST results to pairs of genes between species that were identified as top BLAST hits in each reciprocal BLAST pair (i.e., query sequence for species A returns target gene in species B, query sequence for species B returns target gene species A, query sequence for species A equals target gene in species A, and query sequence for species B equals target gene in species B). Note that this analysis does not preclude a gene in one species pairing with more than one gene in another species. We maintained this possibility in our analysis because differences in genome quality, annotation methods, gene duplication and retention of duplicated genes could result in one gene legitimately being the best high-quality match for more than one gene in another species (and vice versa). In these cases, it can be difficult to determine conclusively which cross-species pair of genes represents the true ortholog. Therefore, we decided to keep all potential orthologous pairs identified by our reciprocal BLAST strategy without requiring 1 to 1 hits; however, we accounted for the possibility of 1 to > 1 hits when determining the number of commonly DE or AS genes across species (i.e., a 1 to 2 reciprocal BLAST result was counted as evidence of a single overlapping pair of orthologs, whereas a 2 to 2 result was counted as two overlapping pairs of orthologs, etc.). Orthologous genes identified by these analyses are available in Table S1.

Splicing patterns of the common AS genes were investigated by plotting the relative exon expression for each gene in each species with a modified version of the plotDEXSeq() function from the *DEXSeq* package.

## Results

### Patterns of differential expression and alternative splicing within species

Overall patterns of differential gene expression in response to cold acclimation in each species were similar to those published previously (Scott and Johnston, 2012; Healy et al., 2017; Metzger and Schulte, 2018), and consequently only a brief description of these results is presented here. In total, 6,719 genes in *F. heteroclitus*, 8,310 genes in *G. aculeatus* and 4,555 genes in *D. rerio* (Table S2) were differentially expressed as a result of cold acclimation. Thus, in all three species acclimation to low temperatures involved changes in expression of a large proportion of the transcriptome (17.9–42.6%). Additionally, substantially more genes increased in expression than decreased in expression in all three species: 68.8%, 65.8% and 61.7% of the total DE genes in killifish, stickleback and zebrafish, respectively (Fig. S1). This pattern had been observed previously for killifish and stickleback (Healy et al., 2017; Metzger and Schulte, 2018), but not for zebrafish (Scott and Johnston, 2012), and the difference between these results for zebrafish is likely due to the use of a modified RNA-seq normalization in the current study. Consistent with this possibility, a bias towards up-regulated gene expression in response to cold acclimation was typical in transcriptomic studies prior to the use of RNA-seq (e.g., Gracey et al., 2004).

In comparison to the substantial proportions of the transcriptome that changed expression quantitatively as a result of cold acclimation, cold-acclimation-mediated qualitative changes in gene expression through alternative splicing were relatively modest. However, acclimation to low temperature still resulted in significant differences in splicing in a large number of genes in each species: 561 in killifish, 426 in stickleback and 866 in zebrafish (Table S2). This likely indicates that the expression of alternatively spliced transcripts is a key component of the responses to chronic exposure to cold temperature in these species. In addition, 51.7%, 66.7% and 41.0% of the AS genes in killifish, stickleback and zebrafish, respectively, were also differentially expressed as a result of cold acclimation (Fig. 1), suggesting that relatively high proportions of alternatively spliced genes are also differentially expressed.

**Figure 1.**
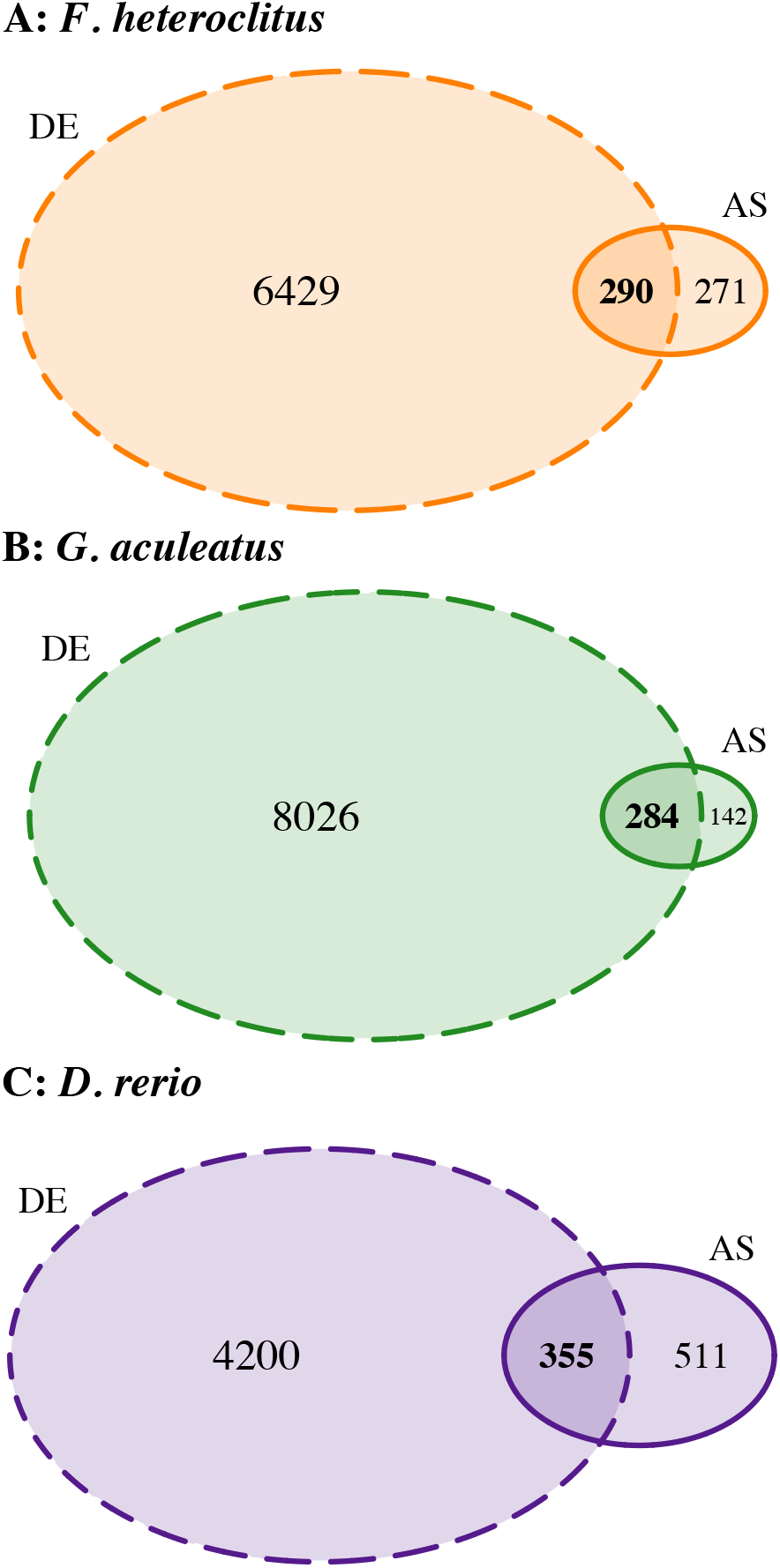
**Venn diagrams displaying the numbers of significant differentially expressed (DE) and alternatively spliced (AS) genes as a result of cold acclimation in *Fundulus heteroclitus* (A, orange), *Gasterosteus aculeatus* (B, green) and *Danio rerio* (C, purple)**. DE genes: dashed outline; AS genes: solid outline.

### Functional enrichment within differential expression or alternatively spliced genes

To summarize potential cellular functions involved in responses to chronic low temperatures we performed GO enrichment analyses for the DE or AS genes in each species. Given the similarities between the DE genes detected in the current study and those observed in previous studies, we expected that GO enrichment analyses would also reveal similar functional enrichments to those published previously (Scott and Johnston, 2012; Healy et al., 2017; Metzger and Schulte, 2018), which they did. As a result, these results are not described in detail here. However, 76, 113 and 29 GO terms were detected as significantly enriched for differential expression in *F. heteroclitus, G. aculeatus* and *D. rerio*, respectively, and lists of these GO terms can be found in Table S3. Importantly in the context of the current study, biological process GO terms for gene expression (GO:0010467), mRNA splicing via spliceosome (GO:0000398) and RNA splicing (GO:0008380), and cellular component terms for catalytic step 2 spliceosome (GO:0071013) and spliceosomal complex (GO:0005681) are among the enriched GO terms with the lowest *p*-values for enrichment in killifish and stickleback (*p* ≤ 9.44 × 10^-9^ for all). Interestingly, these terms are absent from the enriched GO terms in zebrafish, despite the fact that the highest number of AS genes in response to cold acclimation was found in zebrafish (866 versus 561 and 426).

Not surprisingly given the lower numbers of alternatively spliced genes compared to differentially expressed genes (Fig. 1), fewer GO terms were significantly enriched for AS genes regardless of species. There were 27 enrichments in killifish, 15 in sticklebacks and 24 in zebrafish (Table S3). As GO biological processes (rather than cellular components or molecular functions) often provide the most insight into the specific cellular functions demonstrating enrichment, Fig. 2 highlights the significantly enriched biological process terms for all three species. In total across the three species, 25 GO biological processes demonstrated enrichment for alternative splicing. Again, in the context of the current study, one particularly interesting result is that RNA splicing (GO:0008380) was significantly enriched for AS genes in response to chronic low temperature in *F. heteroclitus*. Yet, strikingly, no GO biological process was enriched in all three species, and moreover most terms were enriched in only one of the species. These observations suggest that although many qualitative changes in gene expression as a result of cold acclimation are observed in each species, there is substantial divergence in the cellular processes with excess numbers of alternatively spliced genes. However, there was a clear pattern for functional enrichments of different GO biological processes associated with muscle contraction or regulation of muscle contraction in all three species (e.g., muscle contraction [GO:0006936] in killifish and stickleback, and striated muscle contraction in zebrafish [GO:0006941]), and GO cellular component and molecular function enrichments that are common in all species (e.g., muscle myosin complex [GO:0030018] and structural constituent of muscle [GO:0008307]) also suggest that alternative splicing of genes involved in contraction is likely common across the species. Thus, there may be more conservation of cellular functions associated with cold-acclimation-mediated alternative splicing than is evident based on comparison of specific GO terms alone.

**Figure 2.**
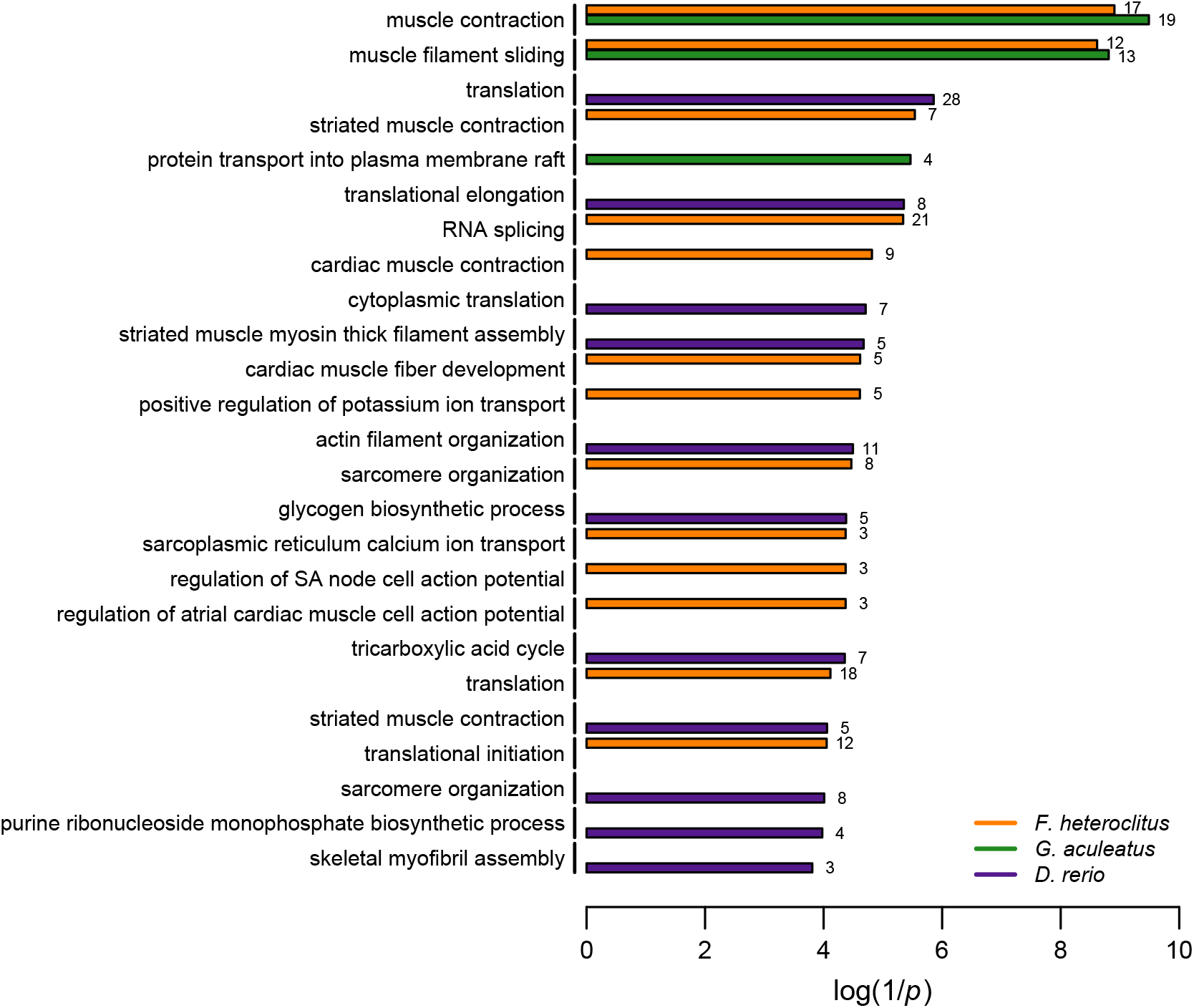
**GO biological processes demonstrating significant enrichment for genes demonstrating alternative splicing in killifish (orange), stickleback (green) and zebrafish (purple)**. Enriched GO terms are listed on the y-axis. Horizontal bars indicate inverse log *p*-values for each term in each species. If a GO term is missing a bar for a species, that term was not significantly enriched in the missing species. Numbers to the right of each bar display the number of alternatively spliced genes in that species with the corresponding GO annotation.

### Comparison of differential expression and alternative splicing across species

To compare patterns of differential gene expression and alternative splicing across killifish, stickleback and zebrafish, we identified putative orthologs across the species for each significant DE or AS gene (see *Materials and Methods*). The overlap in DE or AS genes among species is displayed in Fig. 3. 1,045 common DE genes and 29 common AS genes were detected among the three species (Fig. 3A,B, respectively). Thus, 12.6–22.9% of all DE genes in a species were part of a suite of genes that were differentially expressed in response to cold acclimation regardless of species, whereas 3.3–6.8% of all AS genes were found commonly in all species (percentage ranges are due to differences in total number of DE or AS genes among species). These results reveal a clear pattern that there is substantial divergence among species in both quantitative and qualitative changes in gene expression as a result of cold acclimation. However, the genes that consistently demonstrate differential expression or alternative splicing across species are potentially an important subset of genes that may represent a conserved core response to chronic cold temperatures.

**Figure 3.**
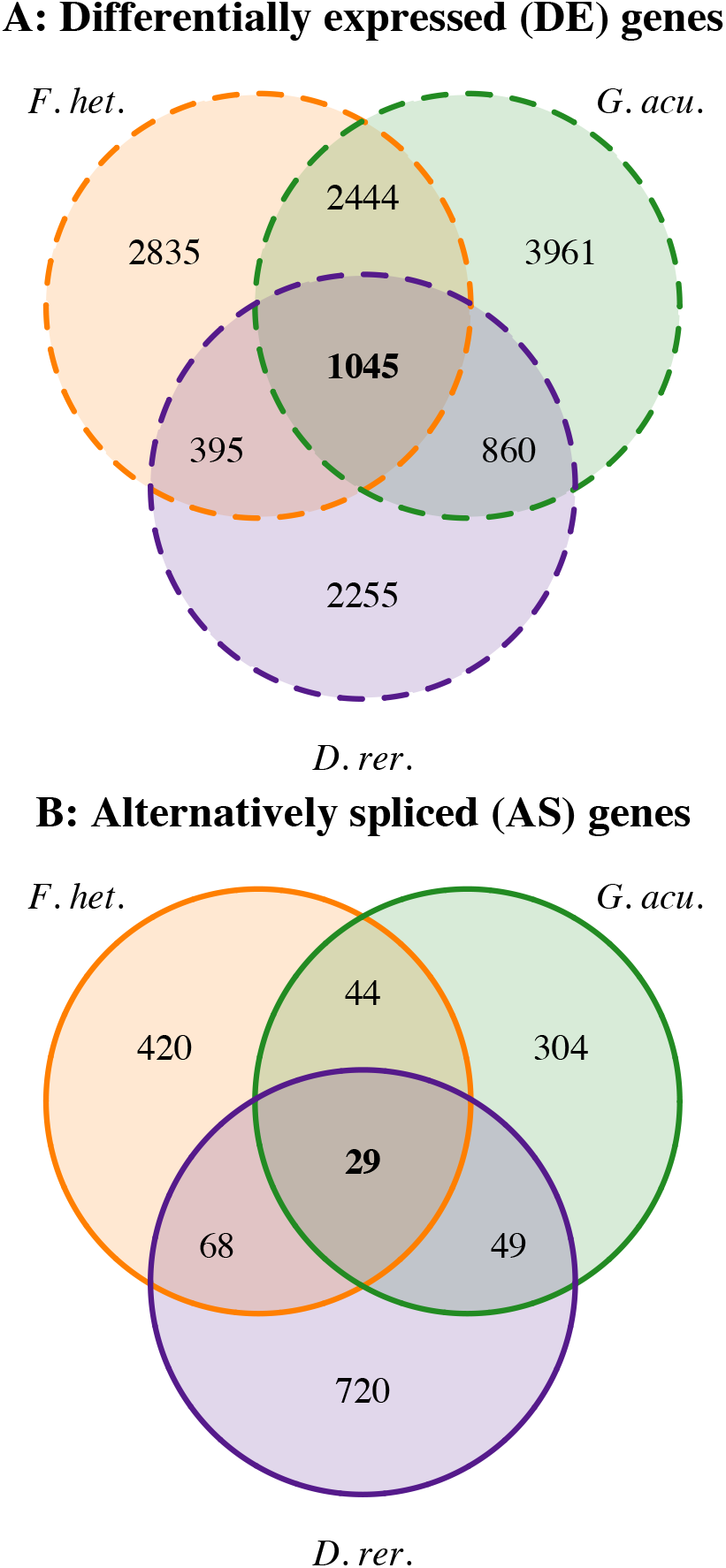
**Venn diagrams displaying the numbers of common differentially expressed (DE; A, dashed outline) or alternatively spliced (AS; B, solid outline) genes as a result of cold acclimation among killifish (orange), stickleback (green) and zebrafish (purple)**. Numbers of DE or AS genes common in all three species are in bold.

The common genes that were differentially expressed as a result of cold acclimation are listed in potential ortholog groups in Table S4. To examine the potential functions that are affected by changes in expression of the common DE genes, we performed GO enrichment analyses for these DE genes in each species. We reduced the influence of differences in GO annotation across the genomes by keeping only the significantly enriched GO terms found in all three species, resulting in 21 significantly enriched GO terms (Table 1). Taken together these GO terms clearly indicate that common DE genes are primarily enriched for involvement in proteasome function (e.g., positive regulation of proteasomal protein catabolic process [GO:1901800]), although there is some indication for enrichment of genes involved in spliceosome function as well (e.g., catalytic step 2 spliceosome [GO:0071013]).

**Table 1.** Significantly enriched gene ontology (GO) annotations for differentially expressed (DE) or alternatively spliced (AS) genes that were identified in all species.

| Common response | GO category | GO term | GO ontology |
| --- | --- | --- | --- |
| DE genes | GO:1901800 | positive regulation of proteasomal protein catabolic process | BP |
|  | GO:0006511 | ubiquitin-dependent protein catabolic process | BP |
|  | GO:0030433 | ubiquitin-dependent ERAD pathway | BP |
|  | GO:0006457 | protein folding | BP |
|  | GO:0000502 | proteasome complex | CC |
|  | GO:0005839 | proteasome core complex | CC |
|  | GO:0019773 | proteasome core complex, alpha-subunit complex | CC |
|  | GO:0031595 | nuclear proteasome complex | CC |
|  | GO:0008540 | proteasome regulatory particle, base subcomplex | CC |
|  | GO:0031597 | cytosolic proteasome complex | CC |
|  | GO:0005634 | nucleus | CC |
|  | GO:0071013 | catalytic step 2 spliceosome | CC |
|  | GO:0005838 | proteasome regulatory particle | CC |
|  | GO:0004298 | threonine-type endopeptidase activity | MF |
|  | GO:0005524 | ATP binding | MF |
|  | GO:0003723 | RNA binding | MF |
|  | GO:0036402 | proteasome-activating ATPase activity | MF |
|  | GO:0016887 | ATPase activity | MF |
|  | GO:0051082 | unfolded protein binding | MF |
|  | GO:0004004 | ATP-dependent RNA helicase activity | MF |
|  | GO:0008307 | structural constituent of muscle | MF |
| AS genes | GO:0006941 | striated muscle contraction | BP |
|  | GO:0005859 | muscle myosin complex | CC |
|  | GO:0030018 | Z disc | CC |

We assessed potential functional enrichments for the common AS genes using the same approach as the one described above for the common DE genes. Three significant GO enrichments were identified for the AS genes observed in all species, which all indicated enriched alternative splicing for genes involved in the structure or function of the contractile apparatus (Table 1). However, with the relatively small number of common AS genes, examining specific genes rather than summary enrichment analyses alone may be more appropriate for these genes. The 29 common AS genes across species are listed in putative orthologous groups in Table 2 (note the number of overlapping genes are calculated as described in the *Materials and Methods*). As expected from the GO enrichment results for these genes, several genes are involved in muscle contraction or the structure of the contractile apparatus (*atp2a1l, myom1b, smyhc1, ryr1, mybph, cacna1s, smyd1b, myh13* and *tnnt2e*). However, genes involved in other cellular functions also demonstrated alternative splicing in all three species. For example, there were cold-acclimation-mediated changes in splicing of genes involved in metabolic processes such as glycogen breakdown (*phkg1* and *phkb*) and glycolysis (*aldoa* and *pfkfb4*).

**Table 2.**
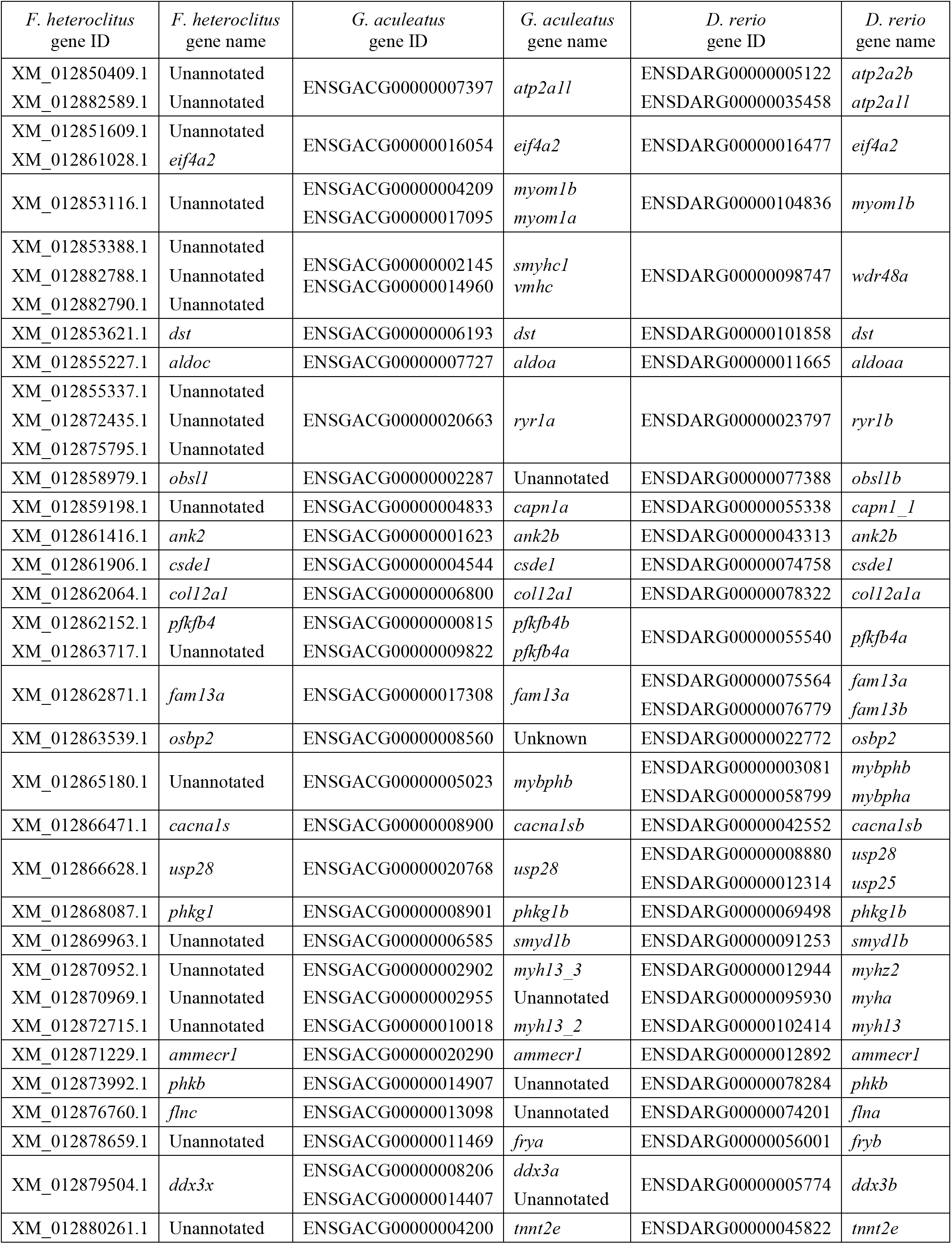
Ortholog groups for the 29 common alternatively spliced (AS) genes identified in all species.

Additionally, cold shock domain-containing protein E1 (*csde1*), which is an RNA binding protein involved in RNA stability and regulation of translation (Mihailovich et al., 2010), is alternatively spliced in all three species. This gene is named for cold shock domains in the protein which were originally identified in bacterial cold shock proteins that increase in expression at low temperature to improve cellular function and growth in the cold (Horn et al., 2007; Mihailovich et al., 2010).

Although there were 29 common AS genes in killifish, stickleback and zebrafish, sharing statistical significance for alternative splicing does not necessarily mean that these genes demonstrate the same splicing patterns in all three species. Indeed, splicing patterns for the common AS genes were generally variable among the species with little evidence for consistent changes in exon usage (*supporting figures will be available as part of an EBI BioStudy should the manuscript be accepted*). However, in many cases there were similarities between at least a pair of the species (e.g., *atp2a1l, eif4a2, fln* and *fry*). Furthermore, two genes had remarkably similar changes in alternative splicing in all three species. For *aldoa* (annotated as *aldoc* in *F. heteroclitus*), an exon near the start of the gene was expressed at higher levels in all of the cold-acclimated fish (*q* ≤ 0.03; Fig. 4A,B,C, note that the orientation of *aldoa* in the *G. aculeatus* genome is in the opposite direction to the orientation in the other two genomes). For *csde1*, an exon or exons in the front half of the gene also had higher usage in fish acclimated to cold temperatures regardless of species (*q* ≤ 0.01; Fig. 4D,E,F). Thus, despite most of the common AS genes sharing only the general characteristic of expression of different splice variants as a result of cold acclimation, two potentially key genes demonstrated conservation of even the specific changes in exon expression across species.

**Figure 4.**
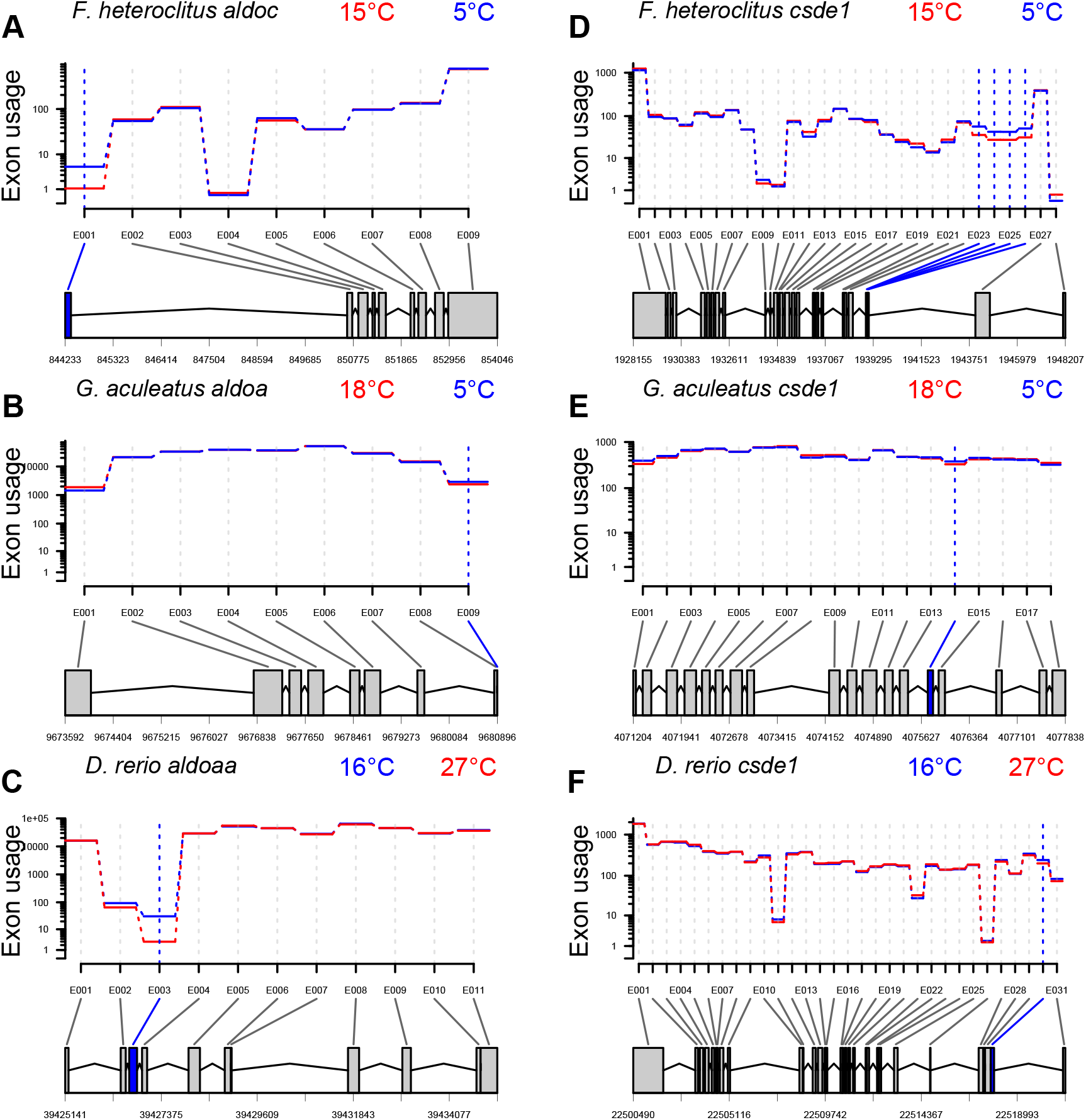
**Plots of relative exon expression (usage) in cold-acclimated (solid blue lines) and warm-acclimated (solid red lines) fish for orthologs of aldolase (A, B, C) and cold shock domain-containing protein E1 (D, E, F) as a result of cold acclimation in killifish (A, D), stickleback (B, E) and zebrafish (C, F).**. Exons with significant differential exon usage are indicated by vertical blue or red dashed lines (blue: higher usage in cold-acclimated fish; red: higher usage in warm-acclimated fish). A graphical summary of exon usage for the flattened gene model for each gene and species is displayed below each plot (grey boxes: exons without differential usage; blue boxes: exons with higher expression in cold-acclimated fish; red boxes: exons with higher expression in warm-acclimated fish; joining lines: introns).

## Discussion

This study clearly demonstrates that qualitative changes in gene expression as a result of changes in mRNA splicing patterns likely play an important mechanistic role in phenotypic plasticity in ectothermic vertebrates. There were substantial changes in splicing patterns in cold-acclimated fish in all three of the species examined in the current study, although fewer genes demonstrated qualitative than quantitative changes in expression. Large proportions of the AS genes were also differentially expressed, and many AS genes were involved in cellular processes that have previously been identified as important components of plastic responses to chronic cold. Furthermore, despite an overall pattern of divergence in cold-acclimation-mediated splicing patterns across species, there was a small set of common AS genes in killifish, stickleback and zebrafish, suggesting that a subset of qualitative changes in gene expression may play a conserved role in regulating plasticity as a result of cold acclimation in ectotherms.

### Role of changes in mRNA splicing patterns in plasticity as a result of cold acclimation

Large-scale quantitative rearrangement of the transcriptome is commonly observed in response to prolonged exposure to cold temperatures in ectotherms (e.g., Gracey et al., 2004), and the results presented here are consistent with this expectation as between ∼20% and ∼40% of the expressed genes in skeletal muscle demonstrated cold-acclimation-mediated differential expression, depending on the species examined (6,719 of 22,120 in *F. heteroclitus*; 8,310 of 19,520 in *G. aculeatus*; 4,555 of 25,477 in *D. rerio*). Generally, these transcriptional rearrangements suggest several hallmark cellular responses to the effects of low temperature: (1) bias towards up-regulation of gene expression, (2) cellular stress responses including up-regulation of genes involved in protein folding, protein ubiquitination, proteasome and DNA repair functions, (3) up-regulation of genes involved in mRNA expression, RNA processing, and pre-mRNA splicing, and (4) enriched differential expression of genes involved in metabolic pathways (Itoi et al., 2003; Gracey et al., 2004; Vornanen et al., 2005; Chou et al., 2008; Castilho et al., 2009; Vergauwen et al., 2010; Long et al., 2012, 2013; Scott and Johnston, 2012; Jayasundara et al., 2013, 2015; Rebl et al., 2013; Bilyk and Cheng, 2014; Mininni et al., 2014; Morris et al., 2014; Wang et al., 2014; Hu et al., 2015; Liang et al., 2015; Verleih et al., 2015; Healy et al., 2017; Ikeda et al., 2017; Metzger and Schulte, 2018). However, the direction of differential expression of metabolic genes is species-specific and likely indicates differences in metabolic strategy (i.e., thermal compensation, no compensation, or inverse compensation) as a result of chronic cold (Precht, 1958; Guderley, 2004; Healy et al., 2017). As expected, the results of our re-analyses to compare cold-acclimation-mediated differential expression across species were consistent with these established patterns. Together, these patterns suggest that the major cellular responses as a result of cold acclimation are thermal compensation of gene expression, up-regulation of protective mechanisms to compensate for the effects of cold on protein and RNA structure and folding, modification of metabolic energy supply and demand, and potentially changes in mRNA splicing patterns.

In comparison to the extent of quantitative changes in gene expression as a result of cold acclimation, qualitative changes to the mRNA pool in response to chronic cold were relatively modest. However, several hundred genes demonstrated cold-acclimation-mediated alternative splicing in each species representing as sizeable transcriptional response. For comparison, quantitative changes in mRNA expression as a result of acute changes in temperature often involve similar numbers of genes as the numbers of AS genes in the current study (e.g., Gasch et al., 2000; Buckley et al., 2006; Schoville et al., 2012; Gleason and Burton, 2015). This suggests that changes in mRNA splicing are key aspects of transcriptional responses to cold acclimation, and that qualitative changes in gene expression are important mechanisms underlying phenotypic plasticity in ectothermic animals.

The possibility that changes in splicing patterns play a key role in cold acclimation is further supported by functional enrichments associated with alternative splicing in killifish, stickleback and zebrafish. Although there was little evidence for common enriched GO terms across the species, which may, in part, be a consequence of differences in the quality or extent of the annotation databases for the different genomes, when the species were considered separately there were functional enrichments that were consistent with several of the major cellular responses to cold acclimation described above. For example, there was enriched alternative splicing for genes involved in RNA splicing (GO:0008380), poly(A) RNA binding (GO:0044822), and the tricarboxylic acid cycle (GO:0006099), suggesting AS genes associated with changes in splice patterns, RNA chaperoning and metabolic pathways in killifish, stickleback and zebrafish, respectively. Similar temperature-mediated changes in exon usage associated with RNA splicing and binding is also common in plants (Capovilla et al., 2015; Filichkin et al., 2015; Calixto et al., 2018). Even at the level of specific genes, variation in splicing patterns was observed for genes, such as cold-inducible RNA binding protein (*cirbp*) in killifish and zebrafish, which have previously been highlighted for their responses and roles in cold acclimation (Gracey et al., 2004; Sano et al., 2015). We also observed substantial evidence for enrichment of AS genes associated with contractile structures and functions, which is clearly consistent with changes in splicing associated with a central function of muscle tissue that is known to demonstrate plasticity in response to temperature change (Johnston et al., 1990; Johnston and Temple, 2002). Therefore, many of the major cellular processes showing plasticity through changes in gene expression as a result of cold acclimation demonstrate not only quantitative but also qualitative responses.

Given the similarities between the cellular processes associated with differential expression and alternative splicing as a result of cold acclimation, it is perhaps not surprising that large proportions of AS genes are also DE genes across the species in the current study (61.7–68.8%). It is possible that this overlap between AS and DE genes is simply a function of the large-scale quantitative transcriptomic changes that are associated with cold acclimation. However, in all three species the percentage of AS genes that are differentially expressed is at least 23.2% higher than the percentage of all expressed genes that are differentially expressed. Thus, it is also possible that our results indicate a bias for differential expression of genes that are also alternatively spliced. Regardless, our data clearly demonstrate that quantitative and qualitative changes in expression are not exclusive at the gene-level, and may highlight genes that play important roles in regulating cold-acclimation-mediated plasticity as there are both changes in transcript amounts and types for these genes. The number of genes demonstrating both quantitative and qualitative responses may also suggest that the functional consequences of the two processes are at least somewhat independent or complementary (i.e., quantitative transcript responses cannot be achieved qualitatively, and vice versa), which would emphasize the importance of understanding the contributions of both types of transcriptomic change in regulating cellular plasticity. With regards to this consideration, an important caveat is the fact that mRNA levels do not always directly reflect protein levels (e.g., Gygi et al., 1999), and the extent to which alternative mRNA transcripts lead to alternative proteins remains unresolved (Tress *et al*., 2017; Blencowe, 2017). Therefore, confirmation that the qualitative changes observed here result in the expression of alternative protein forms with functional differences is an important next step.

One particularly interesting process demonstrating both quantitative and qualitative changes in mRNA expression in the cold is RNA splicing itself. For instance, GO terms associated with RNA processing have enrichment for differential gene expression as a result of cold acclimation in killifish, stickleback and zebrafish, and at least in killifish similar GO terms are also enriched for alternative splicing. Overall thermal compensation of gene expression likely contributes to some of these quantitative changes in the expression of splicing genes. However, qualitative changes in the expression of these genes are consistent with differences in splicing patterns rather than simply the amount of splicing machinery present in the cell. In plants, several genes involved in regulation of splicing express alternatively spliced mRNAs in response to changes in temperature (Lazar and Goodman, 2000; Iida et al., 2004; Palusa et al., 2007), and these effects play a primary role in cellular thermosensory functions (Capovilla et al., 2015). Similar thermosensory functions are associated with temperature-mediated splicing events in yeast (Meyer et al. 2011). Additional work is necessary to conclude that changes in pre-mRNA splicing serve as a cellular thermometer in ectothermic animals as well; however, several splicing factors demonstrate alternative splicing in our study (e.g., *sf3b1, srsf3* and *sfpq*). Thus, there is potentially an under-appreciated and important role for alternative splicing in thermosensory functions in animals, and this possibility merits further research attention in the future (Somero, 2018).

### Conservation of quantitative and qualitative differential gene expression across species

Considering the overall similarities in cellular processes demonstrating differential gene expression and alternative splicing as a result of cold acclimation across studies and species, it is remarkable the extent of divergence in specific differential expression or alternative splicing patterns among even the three species of fish examined in the current study. Moreover, this is the case if either specific genes (DE or AS), or enriched cellular functions associated with these genes are compared. Several possibilities could contribute to these differences in gene expression responses across species. (1) Our reciprocal BLAST strategy for ortholog identification may have been conservative, which would reduce the potential for overlapping DE or AS genes across the species. (2) Metabolic strategies in response to prolonged cold temperatures often differ among species, and previous studies have suggested that killifish do not compensate metabolic functions in skeletal muscle (Healy et al., 2017), whereas stickleback and zebrafish display thermal compensation (Metzger and Schulte, 2018; Scott and Johnston, 2012), which is consistent with divergence of gene expression patterns. (3) Although similar relative cold exposures were compared in our study, the specific temperature ranges for each species may change the effects of cold temperature on cellular function. For example, state-transitional effects of biological membranes occur below 10°C in *F. heteroclitus* which results in dramatic differences in the effects of cold shock at temperatures above or below this threshold (Buhariwalla et al., 2012; Malone et al., 2015). Thus, 15–18°C to 5°C in killifish and stickleback may represent a substantially different physiological challenge than 27°C to 16°C in zebrafish. However, this would suggest that comparisons of the DE or AS genes in killifish and stickleback should be more similar than the same comparisons for either species with zebrafish, and our data are not consistent with this pattern. (4) Lastly, variation in GO annotations (e.g., level of annotation) of the killifish, stickleback and zebrafish genomes almost certainly reduces the overlap in enriched annotations associated with the DE or AS genes in each species, and as discussed above functional overlap is likely higher than comparisons of specific annotations alone. Although these possibilities may partially explain the observed divergence of differential expression and alternative splicing among species in the current study, it is unlikely that they do so completely. Thus, our data suggest that the bulk of either quantitative or qualitative changes in gene expression in response to cold acclimation are not conserved across species.

Despite this overall pattern of divergence in gene expression, there were subsets of both DE and AS genes there demonstrated conservation of quantitative or qualitative changes across killifish, stickleback and zebrafish. These genes may be central in cellular responses to prolonged exposure to cold temperature that occur regardless of species. Consistent with greater numbers of genes demonstrating quantitative changes in gene expression than qualitative changes, higher numbers of DE genes than AS genes demonstrated conserved patterns. Indeed, this exceeded the extent which would be expected from the numbers of DE or AS genes alone, as 12.6–22.9% of all DE genes were commonly identified in all three species, whereas this was the case for only 3.3–6.8% of AS genes. Therefore, our data indicate that there may be more conservation of quantitative than qualitative changes in gene expression as a result of cold acclimation. This raises the intriguing possibility that this may be a common aspect of plastic changes in gene expression in response to other environmental factors as well, and could suggest that qualitative changes in gene expression are a critical in species-specific responses.

Genes that were commonly differentially expressed in response to chronic cold temperature in killifish, stickleback and zebrafish were enriched for those involved in many of the major cellular functions known to respond to cold acclimation described above (e.g., RNA splicing, proteasome function and metabolic pathways). This suggests that at least some aspects of quantitative changes associated with these central cellular responses are conserved across species. In contrast, only functional annotations related to muscle contraction (striated muscle contraction [GO:0006941], muscle myosin complex [GO:0005859] and Z disc [GO:0030018]) were enriched among the AS genes that were commonly identified in the three species. These results may be consistent with differences in the functional consequences of quantitative and qualitative changes in gene expression as suggested above. Given the high structural dependence of muscle contractile functions, structural changes in proteins as a result of changes in splicing patterns may be necessary to adequately adjust contractile function either to reduce energetic demands or to compensate for the slowing thermodynamic effects of low temperatures. This possibility would be consistent with previous work demonstrating temperature-mediated changes in myosin isoform expression in common carp that improve muscle performance at either warm or cold temperatures (Goldspink, 1995). Despite functional enrichment of common AS genes indicating that these genes are not involved in the central responses previously associated with cold-acclimated-mediated changes in gene expression, several common AS genes are known to function in metabolic pathways (e.g., *aldoa, pfkfb4, phkb* and *phkg1*), or RNA binding and chaperoning (e.g., *csde1*). Thus, it is likely that there are conserved changes in pre-mRNA splicing that also contribute to plasticity of cellular processes known to play important roles in cold acclimation. Furthermore, although most of the common AS genes demonstrated variation in patterns of alternative splicing across species, two of these genes (*aldoa* and *csde1*) even had similar changes in exon usage in cold-acclimated fish regardless of species, suggesting that not only are AS genes involved in metabolic pathways and RNA chaperoning, but also that these are key responses with conservation of splicing patterns across species.

## Conclusion

Our results reveal that there are substantial changes in mRNA splicing patterns in response to chronic cold temperatures in ectothermic vertebrates. The extent of these qualitative changes in gene expression is smaller than the extent of quantitative changes in transcript levels overall; however, differences in splicing patterns are observed for genes involved in many of the cellular processes that are thought to be key in cold-acclimation responses. Thus, it is likely that these changes in splicing play an important mechanistic role underlying phenotypic plasticity in response to temperature at the cellular level. In addition, genes demonstrating qualitative changes in expression often also demonstrate quantitative changes in expression, and there are both conserved AS genes and conserved patterns of exon usage among the three species in our study. These observations suggest that there may be changes in splicing patterns for a set of genes that play central roles in cold-acclimation responses across ectotherms. Taken together, our findings emphasize that qualitative changes in gene expression, in addition to quantitative changes, are important physiological mechanisms contributing to phenotypic plasticity as a result of environmental change.

## Data availability

RNA-seq reads utilized in the current study are available at the NCBI SRA (SRP091735 and SRP135801 for killifish and stickleback, respectively), and the EBI Array Express Archive (E-MTAB-1155 for zebrafish). The gene-wise and exon-wise read count files, GO annotation databases, and flattened GFF files for each species will be uploaded to an EBI BioStudy record associated with our study should the manuscript be accepted. Our reformatted GFF file that allowed the creation of the flattened GFF file for *F. heteroclitus* will also be provided with the BioStudy. All other data and results associated with the manuscript are provided as supplemental materials.

## List of abbreviations

Alternatively spliced –: AS
Basic local alignment search tool –: BLAST
Differentially expressed –: DE
European Bioinformatics Institute –: EBI
Gene ontology –: GO
General feature format –: GFF
National Center for Biotechnology Information –: NCBI
Sequence Read Archive –: SRA

